# A Novel and Direct Metamobilome Approach improves the Detection of Larger-sized Circular Elements across Kingdoms

**DOI:** 10.1101/761098

**Authors:** Katrine Skov Alanin, Tue Sparholt Jørgensen, Patrick Browne, Bent Petersen, Leise Riber, Witold Kot, Lars Hestbjerg Hansen

## Abstract

Mobile genetic elements (MGEs) are instrumental in natural prokaryotic genome editing, permitting genome plasticity and allowing microbes to accumulate immense genetic diversity. MGEs include DNA elements such as plasmids, transposons and Insertion Sequences (IS-elements), as well as bacteriophages (phages), and they serve as a vast communal gene pool. These mobile DNA elements represent a human health risk as they can add new traits, such as antibiotic resistance or virulence, to a bacterial strain. Sequencing libraries targeting circular MGEs, referred to as mobilomes, allows the expansion of our current understanding of the mechanisms behind the mobility, prevalence and content of these elements. However, metamobilomes from bacterial communities are not studied to the same extent as metagenomics, partly because of methodological biases arising from multiple displacement amplification (MDA), often used in previous metamobilome publications. In this study, we show that MDA is detrimental to the detection of larger-sized plasmids if small plasmids are present by comparing the abundances of reads mapping to plasmids in a wastewater sample spiked with a mock community of selected plasmids with and without MDA. Furthermore, we show that it is possible to produce samples consisting almost exclusively of circular MGEs and obtain a catalog of larger, complete, circular MGEs from complex samples without the use of MDA.

**Importance:** Mobile genetic elements (MGEs) can transport genetic information between genomes in different bacterial species, adding new traits, potentially generating dangerous multidrug-resistant pathogens. In fact, plasmids and circular MGEs can encode bacterial genetic specializations such as virulence, resistance to metals, antimicrobial compounds, and bacteriophages, as well as the degradation of xenobiotics. For this reason, circular MGEs are crucial to investigate, but they are often missed in metagenomics and ecological studies. In this study, we present, for the first time, an improved method, which reduces the bias towards small MGEs and we demonstrate that this method can unveil larger, complete circular MGEs from complex samples without the use of multiple displacement amplification. This method may result in the detection of larger-sized plasmids that have hitherto remained unnoticed and therefore has the potential to reveal novel accessory genes, acting as possible targets in the development of preventive strategies directed at pathogens.

## Introduction

Microbes have accumulated an immense genetic diversity over time. This genetic variety allows them to live in almost any conceivable environment and is mainly caused by the dissemination of MGEs and genome plasticity, which permits adaptability to environmental stresses (1, 2). MGEs are, in their simplest form, elements of DNA mediating mobility, but these elements are not necessarily incorporated in the chromosome. They encompass integrons, transposons, plasmids, IS-elements, Integrative and Conjugative Elements (ICEs) and bacteriophages (phages) (3, 4). MGEs are known to produce a remarkable impact on genome plasticity and are major contributors to the rapid evolution of bacteria, as horizontal gene transfer (HGT) of MGEs between distant bacterial species can result in the addition of new traits to a strain, such as antibiotic resistance (5). The capability of microbes to take up and express foreign DNA is not limited to antibiotic resistance genes, but includes genes essential for other complete pathways such as nitrogen fixation or the degradation of pesticides or xenobiotics (6–9). This illustrates that MGEs serve as a vast communal gene pool that enable prokaryotes to adapt to stresses and fluctuations in their environments. Such communal gene pools are often referred to as metamobilomes and they constitute a wide range of circular elements such as plasmids, IS-elements, transposons and phages (10). Metamobilomics serves as a powerful tool for the study of accessory genes, commonly carried by MGEs, and the identification of potential therapeutic targets (2, 11). Ghaly and Gillings recently reviewed mobile DNA by comparing it to endoparasites, exploiting bacteria for selfish benefits, and they suggest that it might be more reasonable to treat multi-drug resistant pathogens by targeting the MGEs driving the persistence of antibiotic resistance rather than killing all bacterial species in order to remove the pathogen (12). Additionally, metamobilomics offers the opportunity to study MGEs in transit or with a potential of being horizontally transferred, given that MGEs such as IS elements often have circular topologies or circular intermediates as they detach from the bacterial chromosome. Hence, metamobilomics allows researchers to expand the current understanding of HGT and the movement of MGEs, as well as to identify them in bacterial genomes, discover how widespread they are and how they impact bacterial evolution. The rising incidence of antibiotic resistance exemplifies one of the most relevant threats posed by MGEs, which results in the loss of thousands of human lives annually (13–16).

Analyzing multiple metamobilomes from different natural environments might identify novel accessory genes for potential use in the industry or previously unknown backbone genes, which could be potential targets for preventive strategies against multidrug resistant pathogens, and provide a detailed framework of the mechanisms responsible for DNA mobility. Previously, metamobilome studies have investigated environments such as groundwater (17), wastewater (18), soil (19), rat cecum (14, 20) and cow rumen (21). However, research on MGEs cannot be compared to the extent of work done in metagenomics, despite the impact of MGEs on community structures and evolution. This is due to extensive technical shortcomings such as (i) contamination with chromosomal DNA, (ii) the complexity of assembling short reads stemming from the many repeats within and between plasmids, and (iii) a low incidence of genes encoding functions not directly related to plasmid stability and maintenance (replication, mobilization and toxin anti-toxin (TA) systems) (10, 21). In regards to the limitation of short reads, new sequencing techniques from Pacific Bioscience (PacBio Menlo Park, CA, USA) or third-generation sequencing, such as the Oxford Nanopore Technique (ONT) are able to produce substantially longer reads. This will improve metamobilome assemblies, despite the fact that long reads are not as accurate as the reads obtained from short-read sequencing in general. Moreover, the long-read technologies currently require high-input levels of DNA, but the development in the area more intense than ever (22). A commonly used strategy to sample a metamobilome exploits the shearing of chromosomal DNA into linear fragments during DNA extraction. Circular MGEs are less likely to shear, especially if their length is shorter than approximately 100 kb, and thus remain undigested following the removal of the sheared chromosomal DNA with exonucleases (3, 4, 10). However, to our knowledge, all previous publications using an exonuclease treatment to enrich for circular elements also include a multiple displacement amplification (MDA) step to obtain a sufficient amount of DNA for sequencing as outlined by Jørgensen, Kiil *et al*., (2014) (10). This approach does not yield many plasmid sequences greater than 10 kb in wastewater samples, mainly owing to the MDA step, as demonstrated by Norman *et al.*, (2014) (23). This methodological bias adds a substantial preference towards the amplification of smaller-sized circular DNA elements and results in plasmid-related genes (e.g., mobilization and replication) far outnumbering the diversity of accessory elements (e.g., antibiotic-resistance genes) in most existing metamobilome studies (14, 17, 20, 21, 23). In this study, we show that a direct metamobilome approach minimizes the bias towards enriching small circular MGEs by the MDA step. The MDA step which can effortlessly be avoided for monoculture mobilomes, has never, to our knowledge, been omitted for metamobilomes (10, 24). The advantages of the presented method are evaluated by comparing the abundance of plasmid related reads in a wastewater sample spiked with a mock community of *Escherichia coli* at different steps in the experimental workflow (Fig. 1). This mock community harbors selected plasmids with sizes ranging from 4.3 kb to 52 kb. We document that MDA is detrimental to the detection of larger-sized plasmids, that omitting MDA is feasible and that the improved method presented in this study allows the detection of an unprecedented catalog of large, complete, circular MGE sequences from complex samples.

**Figure 1:**
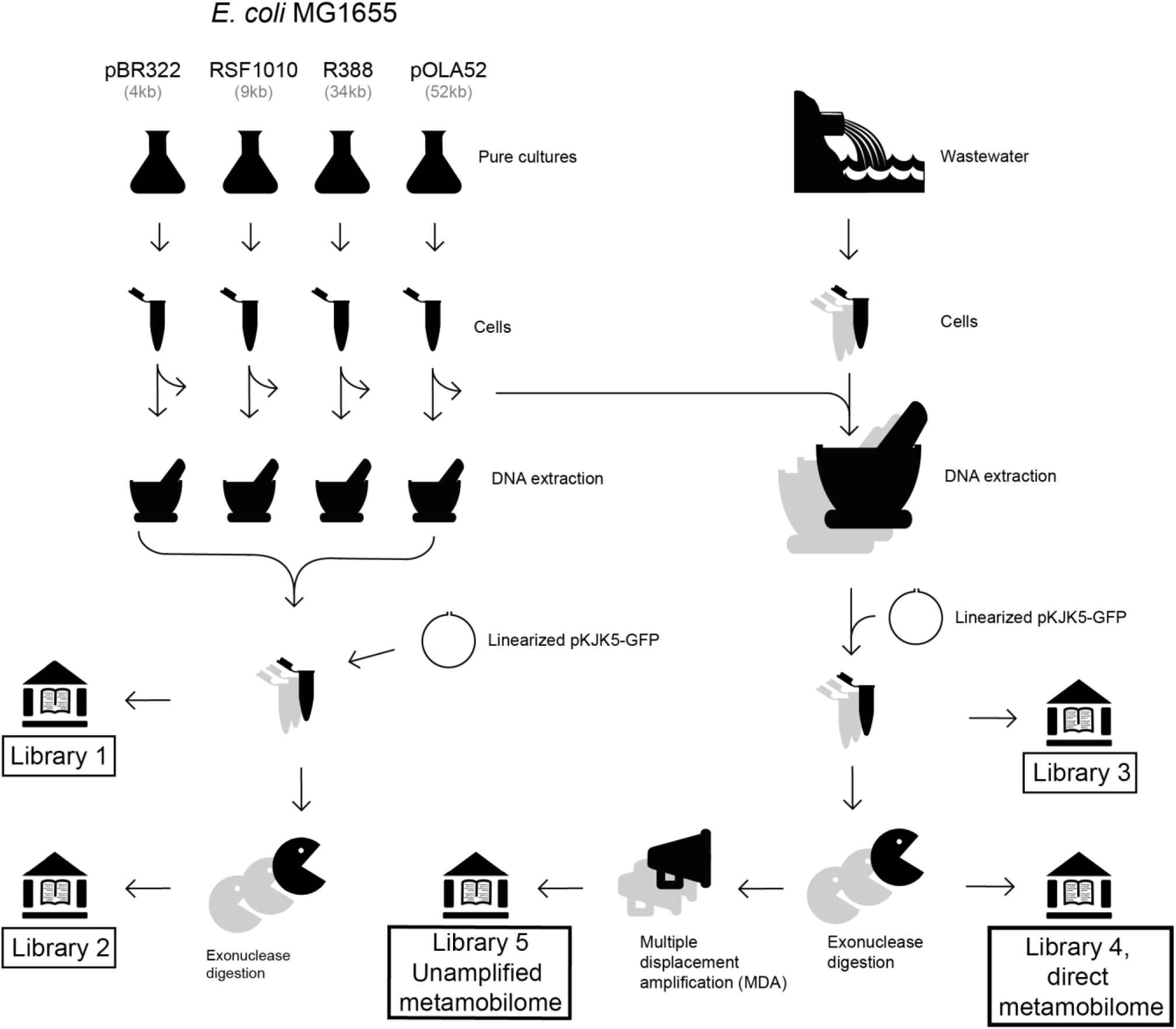
Experimental setup. A total of 5 libraries were constructed in independent triplicate workflows. **Library 1**: Undigested mock community DNA purification, no amplification, and no digestion. **Library 2**: mock community direct DNA metamobilome, no amplification, but digestion. **Library 3**: Undigested wastewater DNA metamobilome with spiked mock community, no digestion and no amplification. **Library 4**: Wastewater **Direct metamobilome** with a spiked mock community, no amplification, but digestion (direct metamobilome). **Library 5**: Wastewater **Amplified metamobilome** with a spiked mock community, digestion, and amplification (amplified metamobilome). Linearized pKJK5_GFP_ was added after DNA extraction from the mock community alone and the wastewater + mock community to monitor the removal of linear DNA. Grey shades indicate the number of replicate samples, which are three replicates for each step. The following creative commons licensed clipart figures were used: http://www.clker.com/clipart-empty-flask-erlen.html,https://openclipart.org/detail/169437/water-pollution,https://www.clker.com/clipart-10885.html

## Results

### Linear DNA is removed by exonuclease treatment

Smaller plasmids (approx. ≤ 100 kb) and other circular extrachromosomal dsDNA elements are not expected to shear during DNA extraction and subsequent processing steps. In order to gauge the efficiency of degradation of linear DNA (chromosomal contamination and, unfortunately, very large circular and linear MGEs) by exonuclease treatment, we determined the relative proportions of reads mapping to four proxies. These were (i) small subunit (SSU) rDNA, hereafter referred to as 16S rRNA, (ii) *Escherichia coli* K-12 MG1655 genomic DNA (25), (iii) the entire sequence of the linearized pKJK5_GFP_ plasmid, and (iv) the GFP gene carried by the linearized pKJK5_GFP_ plasmid. The 16S rRNA of the ribosomal complex was chosen as it has been commonly used for the detection of chromosomal DNA (26). It is considered unlikely for the 16S rRNA to reside on an extrachromosomal element, though this has been observed in some bacteria (27). As the host for all plasmids in the mock community is *E. coli* K-12 MG1655, we find its estimated chromosomal abundances in our datasets a relevant measure in addition to the 16S rRNA. We spiked the wastewater samples with linearized plasmid pKJK5_GFP_ carrying a gene encoding the green fluorescence protein (GFP) because it is extremely unlikely to find *gfp* naturally in wastewater in Denmark, as the gene originates from the Pacific jellyfish species *Aequorea Victoria* (28). Thus, when placed on a plasmid, which is subsequently linearized, it is a very good proxy for the exonuclease degradation of linear DNA. Furthermore, reads mapping to the vector pKJK5_GFP_ itself make a fourth proxy for linear DNA degradation. The vector pKJK5_GFP_ could share genes with elements found in environmental samples, and so may be a less good proxy for linear DNA quantification, but the vectors length of 54 kb makes the resolution much higher than the 965 nt GFP alone (28). The pKJK5_GFP_ and 16S rRNA proxies for linear DNA are somewhat pairwise dependent as *E. coli* K-12 MG1655 harbors 16S rRNA and pKJK5_GFP_ harbors GFP far from the linearization cut site, *Xba*I. As the extraction kit used in this study is designed to enrich the plasmid fraction of a sample, the amount of chromosomal DNA in the untreated DNA purified sample does not directly translate to the amounts of plasmid and chromosomal DNA inside cells. Because of this, we chose to represent the data as a relative measure rather than an absolute measure, by standardizing the value to 100% in samples without exonuclease treatment (Fig. 2).

**Figure 2:**
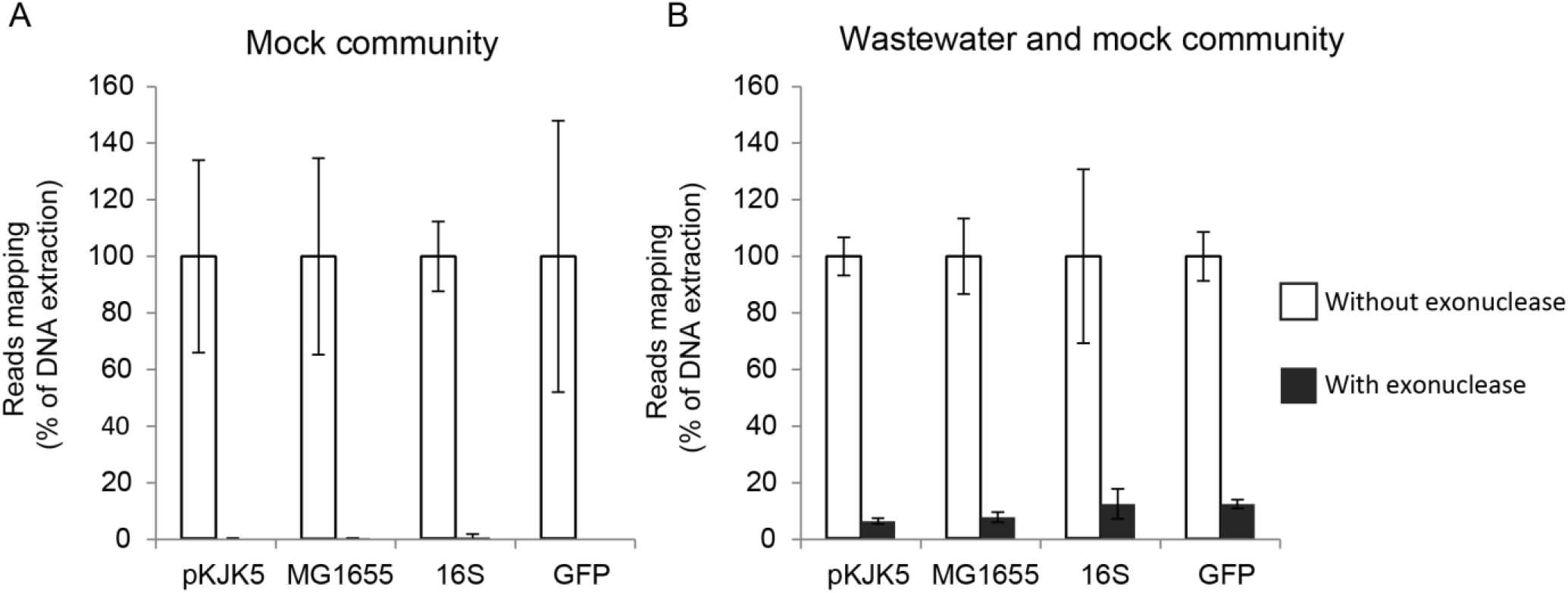
Effectiveness of exonuclease treatment. For each proxy of linear DNA, the average proportions mapping to the proxy in the untreated samples (White columns: Library 1 (A) and Library 3 (B)) were adjusted to 100% using an appropriate multiplication factor. The same factor was used to adjust the average proportion of reads mapping to the proxy after exonuclease treatment (Black columns: Library 2 (A) and Library 4 (B)). Error bars show ± 1 standard deviation about the mean of three replicates.

Results from the mapping of reads from the mock community alone to the complete sequences of its constituents before and after the exonuclease treatment, showed a nearly complete removal of all proxies of linear DNA when normalized to the untreated samples (Fig. 2A). For linear pKJK5_GFP_ and GFP, respectively, an average of 0.38% (± 0.14) and 0% of the relative coverage (relative percentage of reads still mapping) remained after exonuclease treatment. For *E. coli* K-12 MG1655 and 16S rRNA, respectively, an average of 0.48% (± 0.074) and 0.69% (± 1.191) of the relative coverage was left after exonuclease treatment. For all four proxies of linear DNA, significant reductions were thus seen for the mock community, as expected due to the laboratory setup: pure strains, no environmental contaminants and the use of kits optimized for *E. coli*.

Next, we investigated the exonuclease-mediated removal of linear DNA in wastewater samples spiked with the mock community (Fig. 2B). The combined wastewater and mock samples had an average of 6.49% (± 1.01) and 12.58% (± 1.55) of the relative coverage left after exonuclease treatment for pKJK5_GFP_ and GFP, respectively. Similarly, an average of 7.84% (± 1.78) and 12.56% (± 5.32) of the relative coverage was left after exonuclease treatment for *E. coli* K-12 MG1655 and 16S rRNA, respectively.

### Mock community redundancy removal

In order to measure the sequencing depth of each of the mock community plasmids, we mapped the sequencing reads to their reference sequences. To avoid reads originating from one plasmid mapping to other plasmids, we performed an all versus all BLAST search of the sequences of the mock community plasmids and the genomic background (*E. coli* MG1655) in CLCgenomics 8.5.1 (Qiagen Venlo, Netherlands) and removed all sequence regions, which were found to reside on more than one molecule. An overview of the resulting non-redundant database can be seen in Table 1 along with the size and fraction of the non-redundant sequences. For the smaller plasmids, a substantial fraction of sequence was removed in this step. The plasmid pBR322, for example, harbors a beta-lactamase gene and a somewhat similar sequence is found on both the pUC18 plasmid and the MG1655 genome (data not shown). Because all but 2% (53 bp) of pUC18 is covered by pBR322, pOLA52, and the MG1655 genome, we excluded it from further analyses. We confirmed the removal of the redundant sequence by performing a second BLAST search, which showed no redundancy (data not shown). We did not further account for potential end effects of removing redundant sequences, as we do not think it will affect the validity of analysis profoundly. In the mapping analysis, we normalized the number of mapping reads in each replicate to account for the fraction of redundant sequences in each mock community plasmid (Table 1).

**Table 1:** Overview of the genetic background, mock community plasmids and redundancy. Na: not applicable

| Sequence name | Accession number | Size (bp) | Size of non-redundant sequence (bp) | % of non-redundant sequence |
| --- | --- | --- | --- | --- |
| *E. coli* K-12 MG1655 | NC_000913 | 4,641,665 | 4,614,691 | 99 |
| pUC18 | LC129268.1 | 2,686 | 53 | 2 |
| pBR322 | J01749.1 | 4,361 | 1,945 | 45 |
| pOLA52 | NC_010378.1 | 51,602 | 47,501 | 92 |
| R388 | BR000038.1 | 33,913 | 33,913 | 100 |
| RSF1010 | M28829.1 | 8,684 | 8,684 | 100 |
| pKJK5<sub>GFP</sub> | This study | 56,563 | 49,410 | 91 |
| GFP | Na | 965 | 714 | 100 |

### The direct metamobilome method is more sensitive to relatively large circular elements than the amplified metamobilome method

Metagenome sequencing on the NextSeq500 platform yielded a dataset consisting of approximately 86 million direct metamobilome reads from the replicates in library 4, and another dataset consisting of 14 million amplified metamobilome reads from the replicates of library 5 (Table 2). When comparing the reads mapped to mock community plasmids from the direct metamobilome (Library 4) and the amplified metamobilome (Library 5), there is a visible difference where smaller-sized plasmids (pBR322) are relatively more abundant following MDA than larger-sized plasmids (R388 (34kb) and pOLA52 (52kb)), two of which are almost not detected in the MDA amplified metamobilome (Fig. 3A). This mock community serves as a control for the wastewater sample and confirms that bigger plasmids have a relatively higher coverage when using the direct metamobilome method in an environment sample.

**Table 2:** Overview of sample type, sequencing platform used and which figures display the indicated sample data. WW: wastewater sample. MCP: mock community plasmids. Note that for mappings to mock community plasmids, datasets were subsampled to maximum 1M reads. Library workflows are visualized in Figure 1.

|  | Sample type | Data used in Figures |
| --- | --- | --- |
| **Library 1** | MCP, without nuclease digestion | Fig 2A (422K MiSeq reads) |
| **Library 2** | MCP, with nuclease digestion | Fig 2A (463K MiSeq reads) |
| **Library 3** | WW, MCP, without nuclease digestion, no amplification | Fig 2B (3.2M MiSeq reads) |
| **Library 4** | WW, MCP, with nuclease digestion, no amplification | Fig 2B (2.8M MiSeq reads) Fig 3A (2.9M NextSeq reads), Fig 3B (86M NextSeq reads) |
| **Library 5** | WW, MCP, with nuclease digestion and amplification | Fig 3A (2.4M NextSeq reads), Fig 3B (14M NextSeq reads) |

**Figure 3:**
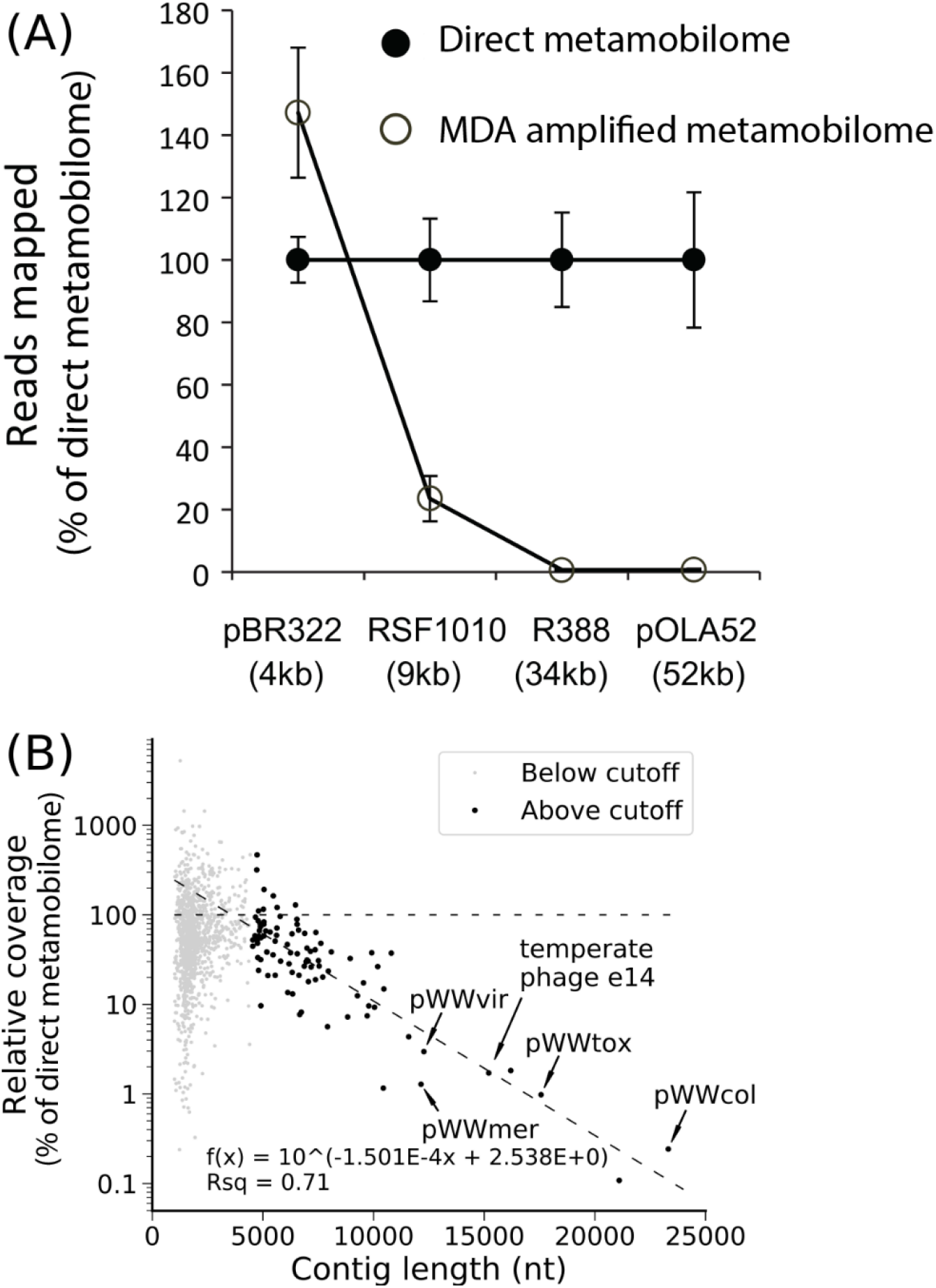
(A) Reads mapped for the mock community plasmids in library 4 and 5. Library 4 is fixed at 100% to visualize the relative differences between amplified and direct metamobilomes. (B) The percentage coverage of the amplified metamobilome (Library 5) relative to the direct metamobilome (Library 4) was plotted against circular element size. The data were divided into two clusters, one with relatively shorter circular elements, (light gray dots; below cutoff of 4 kb), and another with relatively longer circular elements (black dots, above cutoff of 4 kb). The line with long dashes is the trend line fitted to the dark gray cluster. The line with the shorter dashes represents 100 % relative coverage, the level at which the direct and amplified metamobilomes display an equal plasmid coverage. The positions of circular elements detailed in Fig. 4 (pWWtox, pWWmer, pWWvir, pWWcol and temperate phage e14) are indicated with arrows. A total of 52 circular elements smaller than 7 kb could not be represented in this plot due to no coverage in the amplified metamobilome.

Following a meta-assembly, a total of 1,413 circular elements larger than 1 kb were identified from the wastewater sample, with a median size of 1,752 bp, a N50 of 2,153 bp, and a total size of approximately 3.2 Mbp. The largest five, completely assembled, circular elements ranged from 15.2 kb to 23.3 kb. The percent coverage of these circular elements in the amplified metamobilome (Library 5) relative to the direct metamobilome (Library 4) was analyzed in relation to the length of the circular elements (Fig. 3B). The circular elements were divided into two groups using a visually judged cutoff of 4.5 kb. One group contained 1,269 relatively short circular elements and the other group contained 92 relatively longer circular elements. A potential linear trend between relative coverage and fragment length, albeit on a log scale, was visually apparent among the relatively larger circular elements. A trend line was fitted to the cluster of relatively larger circular elements (R^2^ = 0.71) and this trend line crossed 100% relative coverage at around 4 kb. This suggests that the direct metamobilome approach performs better than the amplified metamobilome method for circular elements larger than approx. 4 kb.

### Annotation of selected wastewater plasmids

Using Prodigal for gene prediction followed by HMMscan with the PFAM-A database for a quick scan of plasmid related genes of the 1413 circular elements, we found 59 elements with rep genes, 86 elements containing plasmid recombination genes, 29 elements with mob genes, 46 elements with TA-system genes and 5 elements with T4SS or conjugation related genes when using the PFAM-A database for annotation. A total of 189 elements had at least one plasmid related gene (36 elements had at least two), while 28 elements contained a transposase from an IS-element and only 3 elements contained one or more plasmid related genes together with a transposase (data not shown).

A subset of ten circular elements were selected for complete functional annotation based on a few interesting predicted genes to exemplify the diversity of the plasmids extracted using the direct metamobilome method from the wastewater. The E-values for each protein sequence are from HHpred, unless otherwise stated, and all E-values describe protein similarities. Megablast hits for all 10 full length nucleotide sequences and all conceptually translated protein sequences on the 10 annotated circular elements are presented in supplementary (Supplementary S1 and S2, respectively). Five of the 10 circular elements were visualized and named pWWtox (25 predicted ORFs), pWWmer (13 predicted ORFs), pWWvir (15 predicted ORF), pWWcol (32 ORFs), and temperate prophage e14 (24 predicted ORFs) (Fig. 4). The two plasmids pWWtox and pWWvir contain Rep proteins (RepB E-value=9.8e-29 and KfrA_N E-value=3.7e-21), mobilization proteins (Mob, pWWtox E-value=1.6e-41 and pWWvir E-value=8.8e-25), as well as recombinase proteins (pWWvir E-value=6.3e-30) (Fig 4A and 4E). Both plasmids encode restriction enzyme related proteins (pWWtox; *Eco*RII e-value=4.2e-90 and pWWvir; *Not*I E-value=1.2e-54) and a DNA cytosine methyltransferase (pWWtox; DNA (cytosine-5-) methyltransferase E-value=1.5e-46 and pWWvir; DNA cytosine methyltransferase E-value=3.4e-44), whereas only pWWtox encodes the coupling protein TraD, involved in conjugation (TraD E-value=9.6e-3). Three TA related genes were predicted on pWWtox, and could potentially be one system but none of them had hits to the same system. One has protein similarity to the nucleotidyltransferase AbiEii toxin (E-value=1.2e-13), which is a part of the AbiE phage resistance TA system (29). The gene following AbiEii had results to the RES toxin in the RES-Xre TA complex (E-value=1.3e-19). The last TA related protein had a relatively high E-value to the Phd/YefM family (HHpred E-value=0.11 and Blastp E-value=0.081), hence the simple annotation TA. Two phage related proteins were identified on pWWtox, a bacteriophage replication gene A (GPA) and an integrase (E-value=1.3e-36) (Fig. 4A).

**Figure 4:**
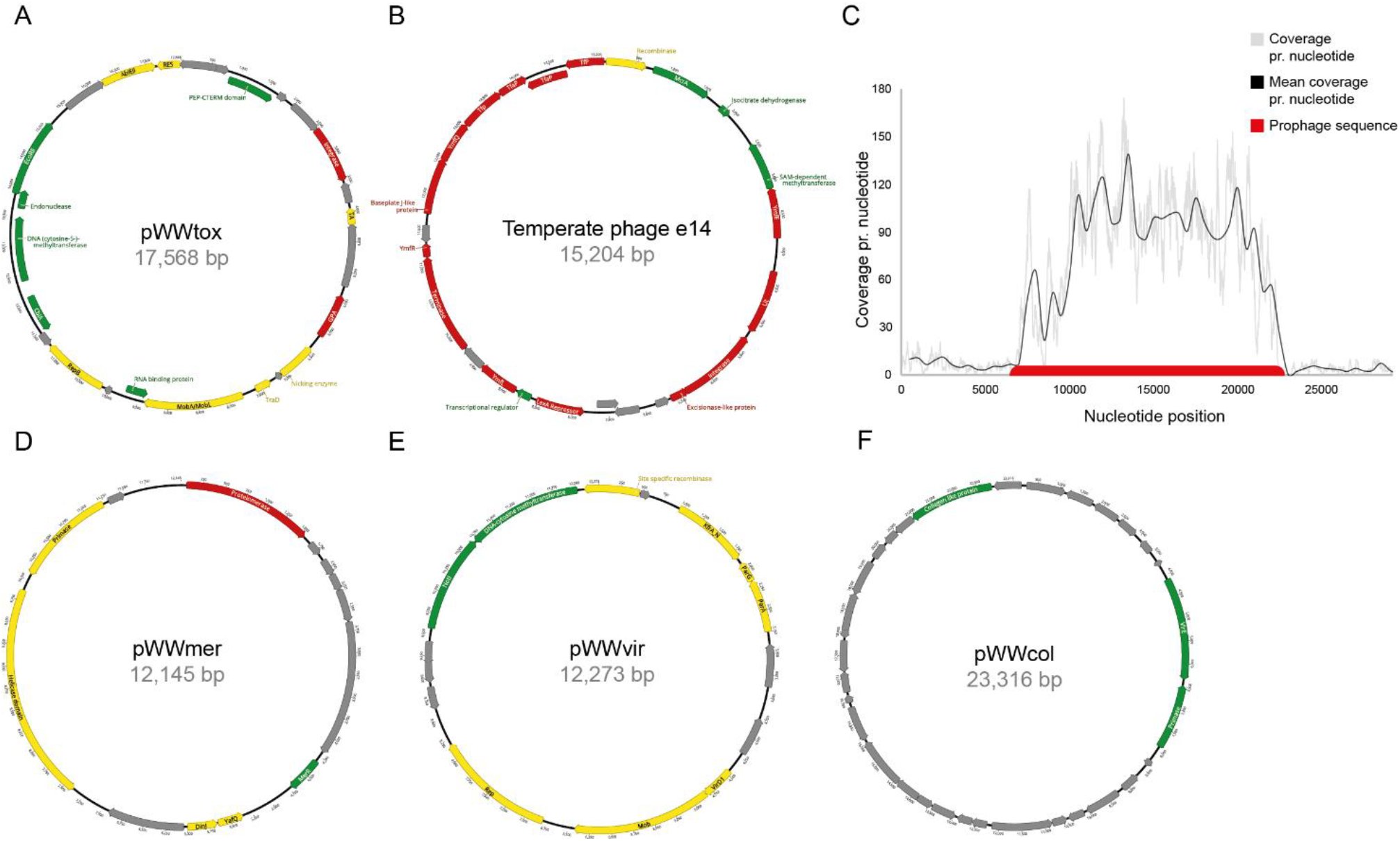
Annotation results of the circular elements pWWtox, pWWmer, pWWvir, pWWcol and the temperate phage e14, together with the read mapping result from the extracted element (B) against the prophage in *E. coli* K-12 MG1655. Grey: hypothetical proteins, Yellow: plasmid related proteins, Red: phage related proteins, Green: Other genes. Each of the four elements were extracted from the direct metamobilome sequence data in Library 4 (Geneious version 11.1 created by Biomatters.(35)).

Among the 10 circular elements selected for annotation, one (temperate phage e14) contained multiple ORFs with similarity to phage-related proteins, including two tail fiber assembly proteins (TfaP E-value=2.6e-18 and E-value=8.2e-21), two tail fiber proteins (TfP E-value=2.1e-7 and E-value=1.2e-20), a terminase (E-value=2.5e-39), a repressor (E-value=3.4e-28) and an integrase (E-value=1.4e-35) (Fig. 4B). Numerous ORFs were denoted as Ymf proteins on the same element (YmfQ, YmfR, YmfL identified by RASTtk), and these are proteins from the cryptic prophage e14 (30). Aligning our circulized element to the cryptic prophage e14 element located in the *E. coli* K-12 MG1655 genome (NC_000913, Table 1), showed perfect consensus. To ensure that the element is a circular element and not a linear genomic residue from the *E. coli* K-12 MG1655 spike-in in our sample, reads were mapped to the prophage element (15,203 bp long) and to flanking regions (7,000 bp up- and downstream) extracted from the full *E. coli* K-12 MG1655 genome (Fig. 4C). This showed highest coverage (16 times higher) across the phage element and lowest coverage at the flanking regions, hence the circular element from the direct metamobilome (library 4) is either the prophage e14 induced from the genome of *E. coli* K-12 MG1655 or from another strain present in the wastewater. Plasmid pWWmer (Fig. 4D) contains the TA-system DinJ-antitoxin and YafQ-toxin (E-value=1.9e-13 DinJ and E-value=4.5e-15 YafQ)(31), as well as a helicase (E-value=3.3e-23), a primase (E-value=2.7e-33) and a phage related protelomerase (E-value= 1.4e-91). However, the most interesting attribute of this plasmid is the MerR protein (E-value=7.9e-16) that regulates the *mer* operon of mercury-resistant bacteria (32, 33). It could be hypothesized that the MerR protein is regulating some of the ORFs annotated as hypothetical, thereby potentially generating a heavy-metal resistance gene cassette similar to the mercury-resistance previously reviewed (32). Plasmid pWWvir contains a gene that encodes a protein with similarity to the VirD1 protein (E-value=1.9e-19), and three hypothetical proteins located *cis* to each other before the endonuclease *Not*I (Fig. 4E). The VirD1 protein is found on the *Agrobacterium* spp. Ti-plasmid (tumor inducing) as reviewed by Gordon and Christie (2014), showing that VirD1 is part of a system that excises transfer-DNA (T-DNA) at its flanking sequences (34). The VirD1 sequence also showed similarity to MobC (E-value=5.1e-6 and 41.8% identity in UniProt) and is potentially a mobilization system together with the adjacent mobilization gene forming the relaxosome mediating conjugation initiation complex. The largest element, pWWcol (23 kb), did not contain any plasmid related functions, but had 29 predicted hypothetical proteins ORFs, a VirE protein, a primase and one collagen-like protein (Fig. 4F). The VirE (E-value=1.1e-11) is a bacterial virulence effector protein related to the Ti-plasmid mentioned previously. The primase (E-value=3.7e-35 hit length 195 bp out of 1428 bp) and collagen-like protein (E-value=8.3e-11 hit length 173 bp out of 1866 bp) are closely related to eukaryotic proteins from *Homo sapiens* and *Rattus norvegicus*, respectively. Blast results from the NCBI and UniProt protein database showed a collagen-like protein too, but originating from a *Clostridium vincentii* (E-value=2e-36 NCBI protein blast) and *Bacillus cereus* (E-value=8.4e-49), respectively. The primase had hits from a *Herpesviridae* (E-value=6.2e-5) too, and the VirE’s second highest hit was to a helicase from a papillomavirus (E-value=4.1e-11). It is not possible to conclude whether pWWcol is viral or a plasmid, especially due to the short hit length. However, these blast hits strongly emphasize the current knowledge gap present in all of the databases, as none of these show a clear result to what this elements is.

## Discussion

### MDA generates a lower frequency of larger-sized elements

While previous studies have indicated that MDA introduces a bias in favor of smaller circular elements (18, 23, 36), the trend has not previously been systematically quantified or shown for environmental circular elements in any mobilomes (10). Here, we use a controlled setup of wastewater and a mock community to show that a MDA step is detrimental to the detection of larger-sized circular sequences. We simultaneously provide an improved direct metamobilome method, which allows for analysis of circular elements previously absent from metamobilome datasets. A direct metamobilome method without amplification will detect a higher diversity of plasmid-encoded genes and theoretically results in a more representative size distribution of detected circular elements. As the study of plasmids is more crucial than ever, e.g., due to the spread of antibiotic resistance, the metamobilomics field needs unbiased research on MGEs. The MDA step is included to produce a sufficient amount of DNA for sequencing. However, due to the nature of the phage rolling circle polymerase Phi29, each Phi29 will amplify small circular elements many times compared to larger circular elements (10, 23, 36). Hence, the coverage of larger elements will be notably lower in a MDA metamobilome, and the natural abundances will be skewed. Our data prove that omitting the MDA step will result in the superior quantification of bigger circular elements.

For this proof of concept, we generated a mock community of selected plasmids (Table 1) with a size range from 4kb to 52kb, and spiked them into a wastewater sample. This ensured that we could detect the relative coverage of larger-sized plasmids (>10 kb) when omitting amplification (Library 4) and compare it to the corresponding relative coverage when including amplification (Library 5). One complication with (but not unique to) the mock community is the repetition of some genes on several plasmids. When these genes are removed in the calculation, the proportions become dissimilar. One such plasmid is pBR322, which only has a proportion of 0.446 unique sequences compared to RSF1010, R388, pOLA52 with fractions of 1, 1 and 0.92053, respectively (Table 1). Despite this redundancy, we documented an obvious diversity of different sized assembled circular elements when omitting MDA and the same result was reproduced in the wastewater sample (Fig. 3). This is the first time the plasmid detection issues in metamobilome studies has been reported systematically, as previous literature on plasmid detection issues has been based on individual samples or has lacked comprehensive analysis of the effect of amplification on the diversity of plasmids (10, 21, 23). The prominent trend in the amplified metamobilome (Library 5) was that larger circular elements were underrepresented or absent, with a related overrepresentation of small circular elements. Comparing the coverage of the amplified metamobilome (Library 5, longest contig in assembly disregarding circularization 14214, and max assembled circle 7kb) with the coverage of the direct metamobilome (Library 4 longest contig in assembly disregarding circularization 49681, and max assembled circle 23.3 kb), the much lower relative coverage for the longer contigs, strongly indicating that the amplified metamobilome has a much lower detection rate for larger plasmids compared to the direct metamobilome (Fig. 3B). These results underline that MDA is detrimental for detection of larger circular elements in the presence of smaller circular elements.

Relatively small circular elements (< 4.5 kb) did not follow the same trend for relative coverage of the direct and amplified metamobilome datasets versus circular element size as seen for larger circular elements (> 4.5 kb) (Fig. 3B). This may be due to false positive identification of relatively small circular elements. One possible explanation of such identification is that complexities, such as *cis*- and *trans*-repeats, in the assembly graph that incorrectly resolve to yield a putative circular sequence are more likely to produce a shorter false positive circular sequence than a longer one. Also, it is probable that redundancy between circular sequences affects the accuracy of the inferred abundances owing to reads mapping randomly given multiple exact matches, with this effect being much more detrimental in the case of smaller circular elements. This was accounted for in the mock community plasmids, by masking redundant elements, but we did not think it feasible to do in the wastewater metamobilome (data not shown). Our analysis indicated that around 4 kb is the size of circular elements above which the direct metamobilome method yields relatively more coverage (Fig. 3B). However, we do not claim this value to be an approximation of where this cutoff would generally lie. From a technical point of view, it would be expected to be influenced by the amount of MDA applied during library preparation. Furthermore, the circular DNA element composition is also likely to affect this threshold. If the abundance of relatively small circular elements in a sample is low relative to that in our spiked wastewater sample, this threshold should appear to be higher and vice versa. Therefore, the value of approx. 4 kb should not be taken as a universal size of plasmid or other circular DNA elements above which it is better to use a direct metamobilome method. Despite expecting that this threshold will not vary considerably, any possible variations have not been examined as this would require extending our experimental design quite considerably. Luo *et al*., (2016) reported a soil metamobilome to potentially contain large plasmids up to 35 kb in size and Jorgensen *et al*., (2017) described circular elements up to 40 kb in a rat gut metamobilome, while Li *et al.*, (2012) detected an abundance of smaller plasmids (<10 kb) in a wastewater metamobilome (18–20). These studies support the notion that the distribution of plasmid sizes might vary between environments or the assembly process varies, but even larger plasmids might have gone unnoticed as all the mentioned studies included MDA in their workflows.

The assembly of circular elements from Illumina reads is, even for direct metamobilome approaches, a challenge. Yet, developments in bioinformatics and sequencing technologies will expand the size range of identifiable plasmids with time. Several programs exist to evaluate the complete, circular plasmid content of a metagenomics sample, including PlacNet, Plasflow, cBAR, Recycler, the HMM+identical contig ends+paired read overlap method used in Jørgensen *et al*,. 2014, and the new and very promising MetaplasmidSPAdes (14, 37–41). After careful consideration, we chose to use Recycler for circularization.

The five largest structures that were successfully circularized from Library 4 ranged from 15.2 to 23.3 kb, and only two contigs were longer (31.8 kb 43.2 kb), but not circularized by Recycler in the assembly graph. However, both contigs had plasmid related genes such as Par, Rep and TA, as well as transposon related genes and could be considered plasmids (data not shown).

### Chromosomal DNA removal

A mobilome can be isolated in several ways, which is discussed in Jørgensen, Kiil, *et al*., (2014), but the most common is to separate all circular elements from the linear DNA by exonuclease digestion (10). This will exclude all plasmids and MGEs displaying linear topologies, as well as very large circular elements that may shear during DNA extraction. It is a limitation of metamobilomics that has not yet been resolved. When isolating circular elements, bacterial chromosomes are expected to shear from their original circular states, even during the gentlest DNA isolation approach. Thus, linearized DNA, such as the plasmid (pKJK5_GFP_) is an acceptable proxy for chromosomal contamination stemming from DNA purification. A recent paper by Kothari *et al.*, (2019) investigated the presence of plasmids in ground water microorganisms, and the authors report retrieval of very large extrachromosomal units (up to 1.7 Mb), but did not sufficiently account for the presence of undigested linearized chromosomal DNA contamination, which could affect the validity of the reported sequences (17). They did use exonuclease treatment of their samples, yet they only removed chromosomal DNA from the assembly by removing contigs with 16S rRNA genes. There are usually 10 to 100 fold more contigs in a given assembly from each genome than there are 16S copies, which is why the majority of chromosomal contigs will not carry 16S rRNA genes (42). Assembling 16S rRNA from metagenomics data is very complicated due to their repetitive nature (43). In comparison to Kothari *et al.*, (2019), we used an ATP dependent exonuclease to remove all linear DNA, but did also quantified the chromosomal content of our samples by using two independent proxies (*E. coli* K-12 MG1655 genome and the vector pKJK5_GFP_). Additionally, we verified the removal of most linear DNA with the same two proxies and an additional two proxies for chromosomal DNA (GFP and 16S rRNA). Reads were mapped to these four proxies in Library 1 and 2 to compare the coverage differences when exonuclease treatment was used or not. It was evident that almost all linear DNA was digested by the exonuclease (Fig. 2A). To ensure that the exonuclease was as effective in the environmental sample, we compared the coverage of Library 3 and 4, which confirmed that the result with added wastewater community was comparable with the mock community alone (Fig. 2B). The 10-fold increase in the proportion of linear DNA could be due to inhibitors in the wastewater decreasing the efficiency of the exonuclease, which would inflate the amount of reads mapping to the linear DNA proxies.

### Annotation of the four plasmids and temperate phage

The annotations show that we get a variety of genes such as heavy-metal resistance *merR*, cobyrinic acid a,c-diamide synthases (*cbiA*) involved in the biosynthesis of cobalamin (vitamin B12) (44), as well as multiple plasmid, phage, transposons and IS-elements related genes. The annotations also emphasize that the databases are not quite mature enough for viral and cryptic plasmid sequences yet, which results in most ORFs being hypothetical (Fig. 4). In addition to the annotation, we observed several reads mapping to eukaryotic 18S rRNA, all belonging to members of the amoebae protist genus *Naegleria*, which interestingly is known to harbor its 18S rRNA genes on a circular, extrachromosomal element of approx. 14kb (45). The temperate e14 phage was, with high certainty, not a genomic DNA contamination from our mock community’s genetic background (*E. coli* K-12 MG1655) as the coverage from our reads across the prophage was much higher than the flanking regions (Fig. 4C). However, we were not able to prove whether the prophage excised from the genome of the added strain or was present in the wastewater community, and prophage induction has been documented previously (24). Nevertheless, this indicates that the direct metamobilome method is capable of uncovering excised prophages, potentially much bigger than e14, therefore has the potential to provide more knowledge about previously unknown or presumably cryptic prophages.

In conclusion, we show here that omitting the Multiple Displacement Amplification step in metamobilome sample preparation will reveal an increased proportion of larger-sized circular elements in a natural sample, which might expand the number of identified and annotated plasmids in the databases. The direct metamobilome approach can, together with advances in long read sequencing and bioinformatics tools, significantly improve the quality of metamobilomics data.

## Material and Methods

### Bacterial strains, plasmids and growth conditions

An *E. coli* MG1655 strain (25) was used as the host to obtain plasmids for the mock community. Plasmids are listed in Table 1. For the construction of plasmid, pKJK5_GFP_, a fragment carrying the p_A1/0403_-gfpmut3-Km^R^ gene cassette was randomly inserted into the plasmid, pKJK5 (accession no. AM261282), using the MuA transposition system (46). The approach was similar to a previous study in which a tetracycline sensitive version of the *gfpmut3* carrying pKJK5 plasmid was constructed (47). Here, cells of *E. coli* GeneHogs transformed with the MuA:p_A1/0403_-gfpmut3-Km^R^ gene fragment were screened for resistance towards kanamycin and tetracycline, but sensitivity towards trimethoprim, using replica plating. The exact location (23.086bp) of the transposon insertion was later identified by sequencing to map 32bp downstream the startcodon of the *dfrA* gene (23.054-23.527bp) encoding trimethoprim resistance. All strains were grown at 37°C, with shaking (250 rpm) for 16h in LB broth medium supplemented with an appropriate antibiotic when needed. Mock community plasmids were isolated using the Plasmid Mini AX kit (A&A Biotechnology, Poland) according to the manufacturer’s instructions.

### Wastewater sample processing and plasmid isolation

Inlet wastewater was obtained from the municipal wastewater plant in Skævinge, Denmark. A total of 1 L of wastewater was used for this study. In order to harvest the microbes, 300 ml of wastewater per sample were centrifuged down for 30 min at 3820 rcf at 4 °C. After harvest, the cells were lysed and plasmid DNA was isolated using the Plasmid Midi AX kit (A&A Biotechnology, Poland) following manufacturer’s instructions. The DNA pellet was dissolved in 500 µl of DNase-free water. DNA was quantified with a Qubit fluorometer using the Qubit^TM^ dsDNA HS Assay kit (Invitrogen, USA).

### Removal of genomic DNA

The wastewater DNA extracts were additionally spiked with a mock community of circular, known plasmids and 5% linearized (linearized with restriction enzyme *Xba*I) pKJK5_GFP_ plasmid (50 ng/ µl) (Table 1, Fig. 2). The Plasmid-Safe ATP-dependent exonuclease (Epicentre, E3101K, USA) kit was used to ensure that mainly circular DNA elements were left in the sample. For each replicate, the exonuclease treatment was set up in a total volume of 50 µl containing: approx. 300 ng of DNA suspended in nuclease-free water, 5 µl of Plasmid-Safe^TM^ 10x buffer, 1 µl of Plasmid-Safe^TM^ exonuclease, 2 µl of 25 mM ATP solution and 2 µl of BSA (10 mg/ml). The mixture was incubated at 37°C for 64 hours. DNA concentration was monitored with a Qubit fluorometer at the beginning of the experiment and after 20, 44 and 64 h of the experiment. After 24 and 48 h, a fresh mixture of 1 µl Plasmid-Safe enzyme and 2 µl of 25mM ATP was added to the mixture. The enriched circular DNA samples were then purified by DNA Clean & Concentrator^TM^-5 kit according to manufacturer’s instructions (Zymo Research, USA). Elution was done with 10 µl of 10 mM Tris (pH 8.0). MDA was carried out as described in (14).

### Sequencing library

The Illumina Nextera XT DNA Library Preparation kit (Illumina, USA) was used for the sequencing library preparation. The DNA input concentration was adjusted to 0.2 ng/μL and the Nextera kit protocol was followed, but 15 cycles were used instead of 12 in the amplification step. The library was stored at −20 degrees before sequencing and analyzed by Qubit fluorometer and PCR for concentration as well as size.

### Read Mapping and coverage of the mock community

Forward reads from each replicate were mapped to the individual non-redundant mock sequences using Bowtie2 v. 2.1.0 with the switches --end-to-end and –sensitive (48). Further, all forward reads were mapped to GFP alone and to the ARB Silva 123 rRNA nr99 small subunit (16S and 18S) ribosomal RNA database. Several chimeric sequences between a potential plasmid/vector and a ribosomal pomegranate sequence were identified, reported to ARB silva, and removed from analysis (data not shown). SAMTOOLS v. 0.1.19 was used for analysis of the mapping result (49). We normalized the fraction of mapping reads in each sample to the mean direct metamobilome treatment counts. Because coverage for each sample and for each plasmid is normalized to the same plasmid, RPKM like normalization is redundant. Replicates with more than 1M reads were subsampled to 1M reads.

### Assembly and circularization

Sequencing reads were quality and adapter trimmed using Trim Galore version 0.4.3 (Brabham Bioinformatics) and assembled using SPAdes 3.12 (50) with the meta switch. The assembly graph was manually curated and only circular paths were retained. All reads were then mapped to the retained putative circular paths using Bowtie2 v. 2.3.4 and the switch --local. Read pairs where at least one of the reads mapped to the putative circular paths were extracted and reassembled using SPAdes 3.12. Circular sequences were then identified using the Recycler pipeline (40). Circular sequences were classified either as plasmids or viruses accordingly (20). Briefly, scaffolds harboring a gene encoding a putative plasmid related gene were classified as plasmid. Sequences with a virus PFAM hit and no plasmid PFAM hit were classified as virus. Sequences with neither were not classified.

### Coverage determination in assembled MGEs

The quality and adapter trimmed read pairs were merged using the-fastq_merge function of usearch v10.0.240 (51) with the arguments “-fastq_maxdiffs 20” and “-fastq_pctid 80”. The merged reads and the paired non-merged reads were mapped separately to the circular sequences identified in the circularization pipeline using bwa mem version 0.7.15-r1140 (52). SAM format read mappings were processed and per-nucleotide read coverages were determined using samtools (depth -a switch) v1.4.1 (49). For each circle, the total coverage was calculated as the sum of the depths for both the merged reads and non-merged reads. A normalization ratio was calculated by dividing the sum of all coverages in the amplified metamobilome by the sum of all coverages in the direct metamobilome. Thereafter, circular sequences with no coverage in either the direct or the amplified metamobilome were omitted from further analysis to avoid zero division errors (52 out of 1413 circular elements discarded). For each remaining circular element, the percentage relative coverage was calculated by dividing the coverage in the amplified metamobilome by the normalization ratio and then dividing that figure by the coverage in the direct metamobilome and multiplying the resulting figure by 100. The percentage relative coverage data along with contig lengths were divided into two clusters using a visually judged cutoff of 4.5 kb. Linear regression was performed for the log-transformed percent relative coverages versus contig length in the cluster containing the longer contigs, which appeared to have a logarithmic linear trend under visual inspection, using the linregress function from scipy package version 1.1.0 in python 2.7.15rc1.

### Annotation of selected plasmids

From all 1413 circular MGEs, ten were picked for annotation and few genes were predicted by Prodigal and roughly annotated by HMMscan with PFAM-A (53). The open reading frame calling and final annotation was done on the selected ten using Glimmer (54), RAST (55), Blast2Go (56), HHpred (57) and PHASTER (58) using default settings, except for RAST which had ‘call-features-insertion-sequences’ enabled. Manual comparisons were performed in Geneious 11.1.5 (35) and CLC Genomic Workbench 11.0 (https://www.qiagenbioinformatics.com/) to verify the coherence of results between all the pipelines. All Open reading frames (ORFs) found automatically were manually curated using the translate tool ExPASy (59) to ensure that reading frames were complete with correct start and stop codons. The ORFs without a correct frame where either removed or amended to be in frame. Genes translated in ExPASy were BLASTed in HHpred (With the following Databases: PDB_mmCIF70_23_Nov, Pfam-A_v32.0, as well as NCBI_Conserved_Domains(CD)_v3.16), NCBI’s BLASTp and UniProt (the latter two with default settings), for further confirmation of the protein homology(60–62). When annotating the function of a predicted gene on a plasmid, the recommendations described elsewhere (63) were followed as closely as possible. Names and function are only given to an ORF if there is consensus in blast results from HHpred, NCBI protein blast and UniProt. The size of the plasmids ranged from 1.6 kb to 23 kb. The three plasmids and one prophage isolated from the wastewater sample were visualized in Geneious 11.1.5 and named with the prefix pWW for plasmid wastewater.

## Acknowledgement

This study was supported by the Human Frontier Science Program (HFSP -RGP0024/2018, KSA & LHH), the Villum Foundation (Block Stipend awarded to L.H.H. and P.D.B. and project AMPHICOP 8960 (TSJ), Lundbeck fund grant no. R44-A4384 (L.H.H. and T.S.J.) and Danish Council for Independent Research (grant no. 4093-00198 awarded to W.K.

