## Supplemental Table 1 for "A Novel and Direct Metamobilome Approach improves the Detection of Larger-sized Circular Elements across Kingdoms"

### Supplementary

S1: Hits for full length sequence blasting in megablast

| Circular element | Hits | E-value | Query cover |
|---|---|---|---|
| RNODE_1224 (1,676 bp) | Transposase for IS481 element from Pseudomonas fluorescens F113 | 0.0 | 84% |
| RNODE_961 (3,425 bp) | Plasmid pSFKW33 form Shewanella sp. 33B | 6e-133 | 12% |
| RNODE_1264 (6,343 bp) | Escherichia coli strain ECONIH4 plasmid pKPC-b33e | 1e-172 | 14% |
| RNODE_1311 (7,919 bp) | Polaromonas sp. H8N plasmid pH8NP1 | 5e-19 | 1% |
| pWWmer (12,145 bp) | Pseudomonas citronellolis strain SJTE-3 (genome) | 0.004 | 1% |
| pWWvir (12,273 bp) | Pseudomonas sp. strain ANT_H54B plasmid pA54BH1 | 1e-168 | 14% |
| Temperate phage e14 (15,204 bp) | Escherichia coli strain K-12 MG1655 | 0.0 | 100% |
| pWWtox (17,568 bp) | Acidiphilium cryptum JF-5 plasmid pACRY04 | 7e-167 | 14% |
| RNODE_1158 (21,098 bp) | Sphingobium chlorophenolicum L-1 plasmid pSPHCH01 | 0.0 | 24% |
| pWWcol (23,316 bp) | Bacillus cereus strain AR156 (genome) | 5e-31 | 1% |
