## Supplemental Table 2 for "A Novel and Direct Metamobilome Approach improves the Detection of Larger-sized Circular Elements across Kingdoms"

### Supplementary

S2: Complete list of all identified Open reading frames (ORFs) for each of the 10 annotated plasmids. ND: not determined

| Contig (circular element) | # identified ORFs | # of hypothetical | Plasmid/Phage/Transposons related proteins | Potential accessory proteins |
|---|---|---|---|---|
| pWWcol | 32 | 29 | ND | Collagen like protein, Primase, VirE |
| RNODE_1158 | 26 | 16 | Recombinase, Type IV secretory system Conjugative DNA transfer, TraD/Mobilization protein, MobA/MobL, Putative toxin VapC6, Recombinase x2, ParA, and RepB. Tn3 family transposase | Thermonuclease |
| pWWtox | 25 | 10 | Site-specific integrase, Toxin-antitoxin x2, AbiEii, Nicking enzyme, TraD, MobA/MobL, and RepB. | PEP-CTERM domain, CbiA, DNA (cytosine-5-)methyltransferase, endonuclease, EcoRII |
| Temperate phage e14 | 29 | 5 | Recombinase, YmfE, Lit, Integrase, Excisionase-like proteins, LexA repressor, YmfL, Terminase, YmfR, Baseplate J-like protein, YmfQ, Tfp x2 and Tfap x2 | mcrA, Isocitrate dehydrogenase, SAM-dependent methyltransferase |
| pWWvir | 15 | 6 | KfrA_N, ParG, ParA, VirD1, Mob, Rep, and Site-specific recombinase | NotI and DNA-cytosine methyltransferase |
| pWWmer | 13 | 7 | Protelomerase. YafQ toxin. RelB antitoxin, Helicase domaine, and DNA primase | merR. |
| RNODE_1311 | 8 | 3 | DNaB replicative DNA helicase, Toxin-antitoxin, Recombinase, ParA, and DNA primase. | ND |
| RNODE_1264 | 6 | 1 | KfrA_N, MobC, RepB, RepA, and RepC. | ND |
| RNODE_961 | 7 | 5 | Replication initiator protein | Integrase |
| RNODE_1224 | 3 | 1 | IS481 family transposase | ND |
